# Accelerated cryo-EM structure determination with parallelisation using GPUs in RELION-2

**DOI:** 10.1101/059717

**Authors:** Dari Kimanius, Björn O Forsberg, Sjors HW Scheres, Erik Lindahl

**Affiliations:** Department of Biochemistry and Biophysics, Science for Life Laboratory, Stockholm University, Sweden; MRC Laboratory of Molecular Biology, United Kingdom; Swedish e-Science Research Center, KTH Royal Institute of Technology, Stockholm, Sweden

**Author notes:** These authors contributed equally to this work.

## Abstract

By reaching near-atomic resolution for a wide range of specimens, single-particle cryo-EM structure determination is transforming structural biology. However, the necessary calculations come at increased computational costs, introducing a bottleneck that is currently limiting throughput and the development of new methods. Here, we present an implementation of the RELION image processing software that uses graphics processors (GPUs) to address the most computationally intensive steps of its cryo-EM structure determination workflow. Both image classification and high-resolution refinement have been accelerated up to 40-fold, and template-based particle selection has been accelerated almost 1000-fold on desktop hardware. Memory requirements on GPUs have been reduced to fit widely available hardware, and we show that the use of single precision arithmetic does not adversely affect results. This enables high-resolution cryo-EM structure determination in a matter of days on a single workstation.

## Introduction

With the advent of direct-electron detectors and advanced methods of image processing, structural characterisation of macromolecular complexes to near-atomic resolution is now feasible using single-particle electron cryo-microscopy (cryo-EM) (Li et al., 2013; Bai et al., 2013). Although this has caused a rapid gain in its popularity, two technological factors still limit wide applicability of cryo-EM as a standard tool for structural biology.

First, partly due to the steep increase in demand, access to high-end microscopes is limited. This is being addressed with acquisition of new equipment in a large number of departments worldwide, as well as the establishment of shared infrastructures (Saibil et al., 2015). Second, processing the increasing amounts of data produced by these microscopes requires computational hardware that is not directly accessible to many labs. Even at larger centres the computational requirements are so high that cryo-EM now suffers from a computational bottleneck. The work presented here addresses this second problem, to the end of drastically reducing the computational time and investment necessary for cryo-EM structure determination.

A typical cryo-EM data set may constitute hundreds or thousands of images (called micrographs) of a thin layer of vitreous ice in which multiple individual macromolecular complexes (called particles) are imaged. Because radiation damage imposes strict limitations on the electron exposure, micrographs are extremely noisy. Thus, to extract fine structural details, one needs to average over multiple images of identical complexes to cancel noise sufficiently. This is achieved by isolating two-dimensional particle-projections in the micrographs, which can then be recombined into a three-dimensional structure (Cheng et al., 2015). The latter requires the estimation of the relative orientations of all particles, which can be found using a wide range of different image processing programs like SPIDER (Frank et al., 1981), IMAGIC (van Heel et al., 1996), EMAN2 (Tang et al., 2007), SPARX (Hohn et al., 2007), FREALIGN (Grigorieff, 2007), XMIPP (Scheres et al., 2008) and RELION (Scheres, 2012a).

These programs also need to tackle the problem that any one data set typically comprises images of multiple different structures; purified protein samples are e.g. rarely free from all contaminants. Multiple conformations, non-stoichiometric complex formation, or sample degradation are all possible sources of additional data heterogeneity. The classification of heterogeneous data into homogeneous subsets has therefore proven critical for high-resolution structure determination and provides a tool for structural analysis of dynamic systems. However, identifying structurally homogeneous subsets in the data by image classification algorithms adds computational complexity, and often increases the computational load dramatically.

An increasingly popular choice for processing cryo-EM data is an empirical Bayesian approach to single-particle analysis (Scheres, 2012b) implemented in the computer program RELION (Scheres, 2012a). In the Bayesian framework, optimal weights for all orientations and class assignments, as well as the three-dimensional reconstruction itself, are learnt from the data in an iterative manner. This allows high-resolution structure determination with minimal bias or user input. In addition, the Bayesian approach has proven highly effective in classifying a wide range of structural variation, such as conformational dynamics within protein domains (Bai et al., 2016), or of very small sub-populations in large data-sets (Fernández et al., 2013). Unfortunately, the regularised likelihood optimisation algorithm that underlies these calculations is computationally demanding. We estimate that a recent 3.7 Å structure of a yeast spliceosomal complex (Nguyen et al., 2016) required more than half a million CPU hours of classification and high-resolution refinement. Computations of this magnitude require the use of high-performance computing clusters with dedicated staff, and restricts development of new algorithms and methods which could benefit the field.

One of the most important recent developments for other scientific programs has been the introduction of hardware accelerators, such as graphics processors (GPUs). To exploit this type of hardware, substantial redesigns of algorithms are required to make many independent tasks simultaneously available for computation, which is known as exposing (low-level) parallelism. However, the possible gain is equally substantial; together with commodity hardware it has been a revolution e.g. for molecular dynamics simulations (Salomon-Ferrer et al., 2013; Abraham et al., 2015), quantum chemistry (Ufimtsev and Martinez, 2008), and machine learning (Jia et al., 2014). Historically, RELION has scaled to the large resources it needs by utilizing image-level parallelism to subdivide the computational tasks within each iterative refinement step, like that of most available alternatives (Fernandez, 2008). However, lower-level core computations on single images in RELION have remained serialised since its introduction more than four years ago.

Here, we describe a new implementation of the regularised likelihood optimisation algorithm in RELION, using GPUs to address its computational bottlenecks. We have chosen to implement our increased parallelism in CUDA, a programming language provided by NVIDIA. The CUDA language currently dominates the GPU computing market, and provides a stable programming environment with a rich C++ interface. We also utilise a number of libraries provided within the CUDA framework, such as cuFFT for fast Fourier transforms (FFTs), and CUB/thrust for sorting and other standard functions. In addition to high-end professional cards there is wide availability of cheap consumer hardware that supports CUDA, which provides outstanding value for many research groups. However, the acceleration and parallelization approaches are general and should be possible to port to other architectures in the future.

The present acceleration of RELION addresses the most computationally intensive steps in a typical image processing workflow. This includes classification of data into structurally homogeneous subsets (2D or 3D classification) (Scheres, 2012b); high-resolution refinement of each homogeneous such set of particles (3D auto-refine) (Scheres, 2012a); and the alignment of movie frames from fast direct-electron cameras (movie refinement) (Scheres, 2014). In addition, we describe an improved algorithm for semi-automated selection of particles from micrographs (Scheres, 2015), this too targeting GPUs. Memory requirements have been reduced to fit widely available consumer graphics cards, and we show that the current adaptation to use single precision floating-point arithmetic does not cause loss of resolvable detail in relion. These developments enable high-resolution cryo-EM structure determination in a matter of days on individual workstations rather than relying on large clusters, and will make it possible to pursue new algorithms for classification and data processing that were previously too expensive even on supercomputers.

## 2 Results

### 2.1 Acceleration of regularised likelihood optimisation

#### Parallelism in the algorithm

The most demanding computation in regularised likelihood optimisation is the comparison of projections of the reference structure along many different orientations with thousands or millions of individual particle images (Fig. 1). This task has several inherent levels of parallelism that can be executed simultaneously. In our implementation, classes, as well as image translations and orientations within each class, are all treated as independent tasks that are scheduled independently on GPUs. Even individual image pixels are evaluated independently of one another (Fig. 2). Reformulation of internal algorithms for such massively parallel streams of data is key to perform well on highly parallel hardware like GPUs. To permit this parallelism, all data required by all tasks must be available before any task can start. The straightforward solution is to prepare all the necessary data prior to starting GPU execution. However, pre-calculating reference projections for all examined orientations and classes in this way requires both very large and fast memory. In practice, this imposes severe limitations on the number of simultaneously examined classes that can be used. To overcome this, our implementation instead stores oversampled Fourier transforms of every class-reference in GPU memory, and extracts 2D-slices (in any orientation) on demand. By utilising fast-access data structures known as textures (normally used to project images on 3D objects), on-demand projection in fact achieves faster execution compared to reading pre-calculated projections from memory. As indicated, this also improves scaling with respect to the number of examined orientations and classes, both in terms of memory utilisation and execution speed. In fact, using GPU-enabled relion, the time needed for classification is only weakly dependent on the number of classes used, whereas the CPU-based implementation has a much steeper linear dependence (Fig. 3).

**Figure 1:**
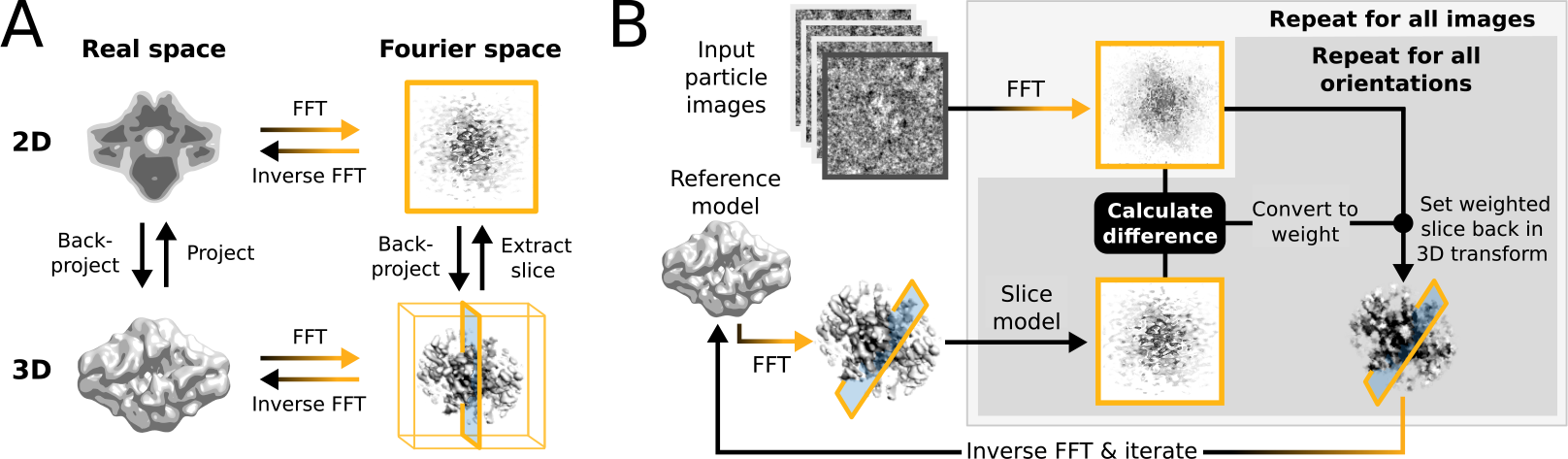
*(A) Operations and the real/Fourier spaces used during (B) image refinement in* RELION. *Micrograph input and model setup use the CPU. Most subsequent processing steps have been adapted for accelerator hardware. The highlighted orientation-dependent difference calculation is by far the most demanding task, and fully accelerated. Taking 2D slices out of (and setting them back into) the reference transforms has also been accelerated at high gain. The inverse FFT operation has not yet been accelerated, but uses the CPU.*

**Figure 2:**
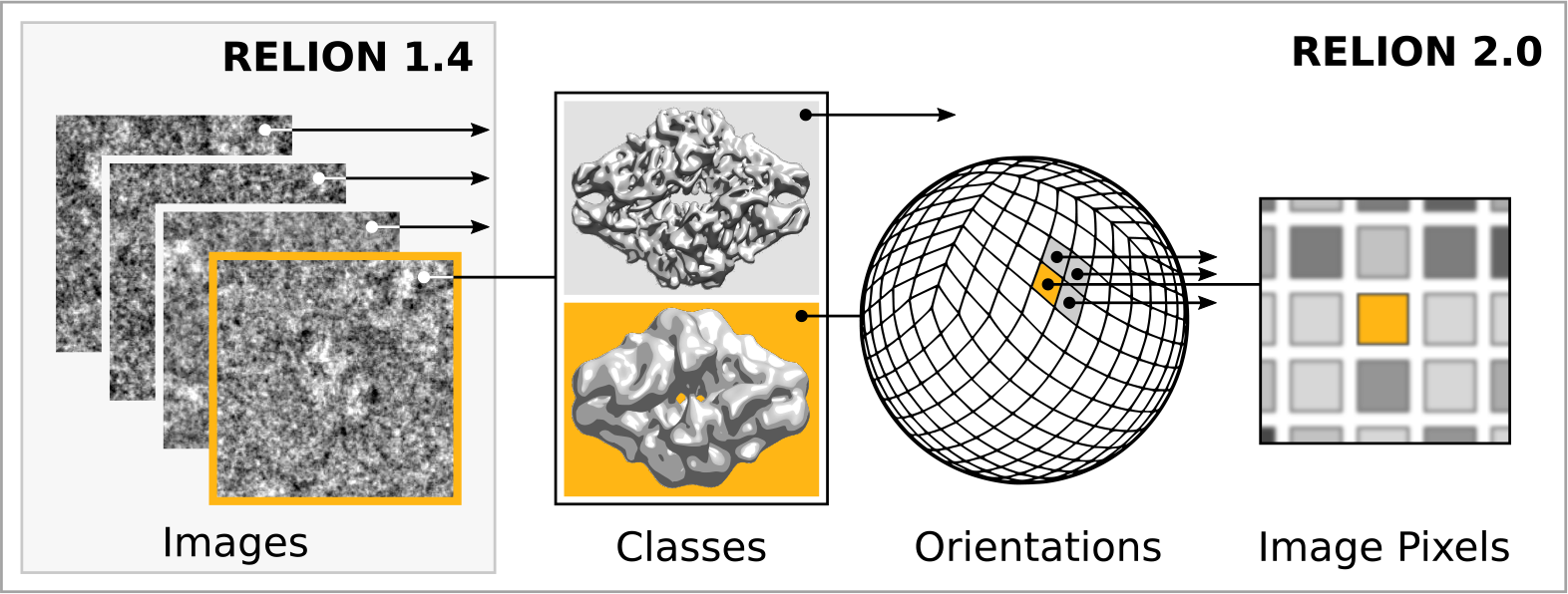
*Extensive task-level parallelism for accelerators. While* RELION-*1.4 only exploited parallelism over images (left), in the new implementation classes and all orientations of each class are expressed as tasks that can be scheduled independently on the accelerator hardware (e.g. GPUs). Even individual pixels for each orientation are calculated in parallel, making the algorithm highly suited for GPUs.*

The result of these adaptations is an implementation that is efficient enough to reliably run on workstations with a single consumer-level GPU, but that also scales favourably to multiple cards, multi-node hardware, and large GPU-based computer clusters. Neither method nor behaviour of RELION has changed from that of version 1.4.

#### Performance

The performance of our implementation on a workstation equipped with modern GPUs can exceed that of hundreds of CPU cores (Fig. 3). This is most prominent for increasing numbers of pixels, orientations and classes, due to the increased low-level parallelism relion-2 has been designed to use efficiently. Therefore, calculations where many classes and orientations need to be sampled, e.g. 3D-classifications over multiple classes and with high orientational sampling rates, experience the greatest gain from the current work (Fig. 3D). In the CPU-only version, the computational time scales linearly with increased number of classes (Fig. 3B) due to the serialised single-image calculations, whereas GPU-enabled execution can show better-than-linear scaling. This is due to the added parallelism and subsequent possibility of concurrent execution, resulting in latencies being hidden. The small parts of the calculation that have not yet been GPU-accelerated provide the largest part of this scaling component (Fig. 3C), which indicates new bottlenecks are now limiting scaling behaviour.

#### Limited precision & accuracy

Floating-point computations inherently lose precision during calculations, due to their finite resolution, which can lead to accuracy problems. RELION has used so-called double precision since its first release, in order to retain as much available accuracy as possible. This is common for scientific programs, which generally have quite stringent demands on numerical results compared to e.g. visualisation applications. While there are professional GPUs with good double precision performance, the consumer market is dominated by visualisation and game applications, and for this reason cheap hardware only provides good performance for single precision. Even for professional hardware single precision improves performance, although the difference is smaller. This makes it highly desirable to use single precision arithmetics wherever possible. In addition to much better floating-point throughput, single precision calculations reduce the memory requirements by 50%, and modern GPUs provide special hardware features for more advanced operations in lower precision. An example of this is the resampling of image rotations, where target pixels are calculated even when not directly overlapping with source-pixels. RELION has always used linear interpolation from proximal pixels, and now does so by utilising special data storage and hardware interpolation on GPUs (so-called textures). Because the required precision depends on the algorithms used in the application, part of the development of relion-2 was to evaluate image refinement quality when using single precision. While we indeed observed a slight loss of precision e.g. in fast interpolation intrinsics, the orientational probabilites of images did not display differences that causes subsequent alteration to the final reconstruction. Execution of the iterative gridding algorithm that underlies the reconstruction step in RELION (Scheres, 2012a) however appeared to show significant loss of information. Therefore, we opted for a hybrid implementation of the algorithm. In this version, the demanding slice projection and probability calculations are performed in single precision on the GPU, while the reconstruction step remains in double precision on the CPU. With this implementation, the probability distributions that result from the difference calculations between reference projections and experimental images do not exhibit any loss of information (Fig. 4).

**Figure 3:**
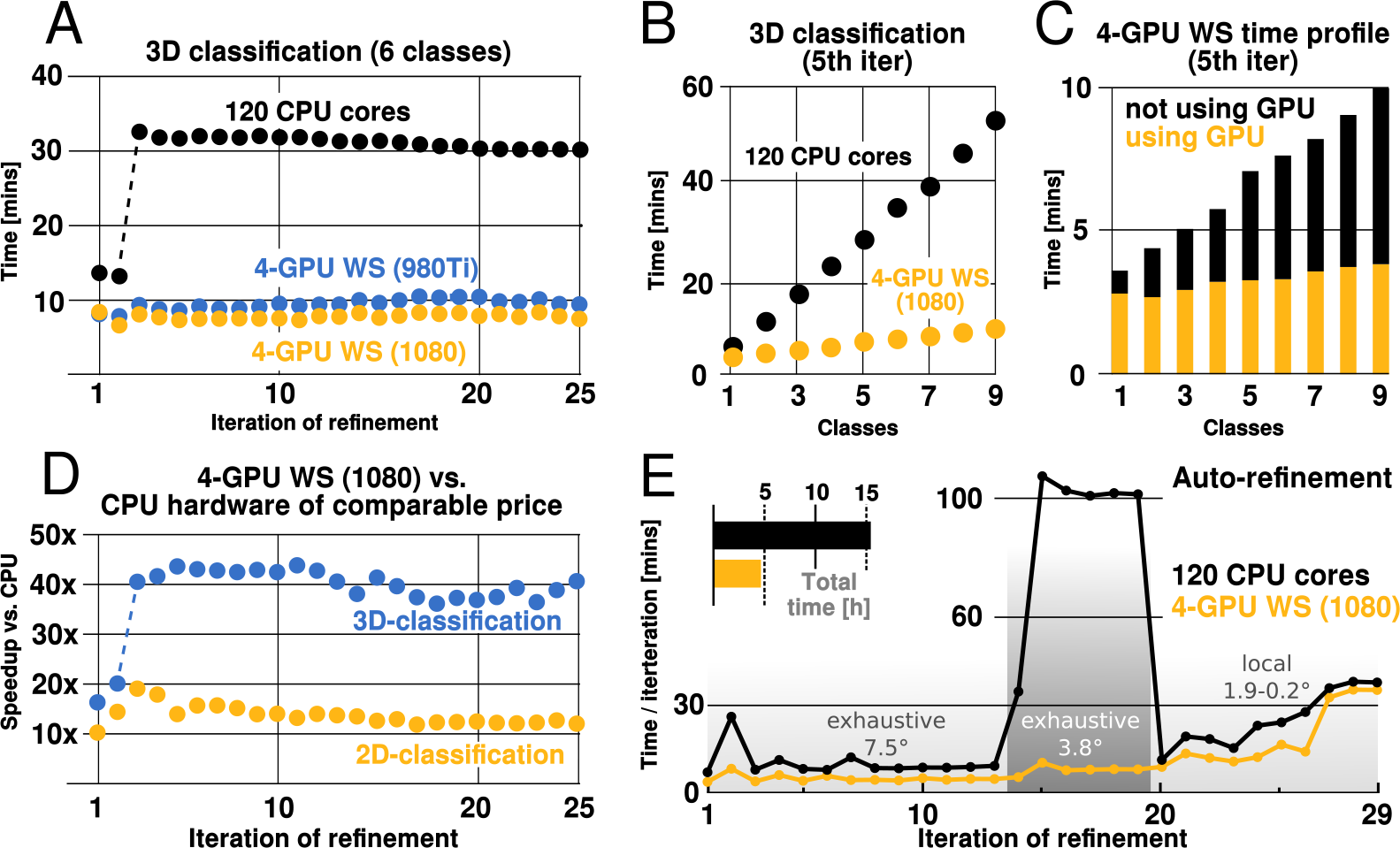
RELION-*2 enables desktop classification and refinement using GPUs.* EMPIAR *(Iudin et al., 2016) entry 10028 was used to assess performance, using refinements of 105k ribosomal particles in 360^2^-pixel images. (A) A workstation equipped with four GPUs easily outperforms even a large cluster job in 3D classification. (B) Additional classes are processed at negligible cost compared to CPU-only execution, due to faster execution and increased capacity for latency hiding. (C) With increasing number of classes, the time spent in non-accelerated vs accelerated execution increases. (D) Comparing a single CPU cluster node to a GPU workstation of roughly comparable price, speedup ranges from an order of magnitude (2D classification) up to a factor 40 (3D-classification using 6 classes). (E) The same workstation also clearly beats the cluster job for single-class refinement to high resolution, despite the generally lower degree of parallelism. During finer exhaustive sampling of the orientational space, the timing difference is even more conspicuous due to the GPU’s ability to parallelise the drastically increased number of tasks.*

**Figure 4:**
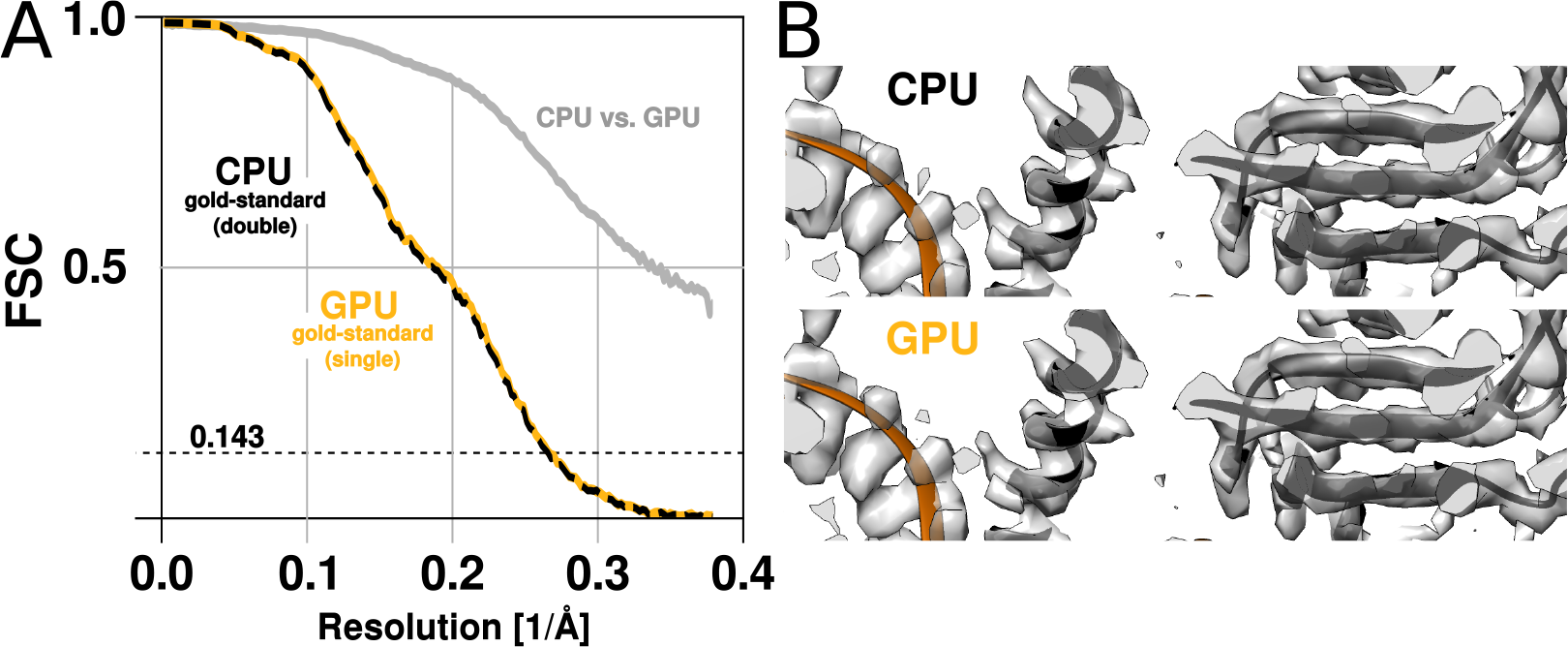
*The GPU and CPU implementations yield qualitatively identical results. (A) A high-resolution refinement of the Plasmodium falciparum 80S ribosome using single precision GPU arithmetic achieves a gold-standard Fourier shell correlation (FSC) indistinguishable from double precision CPU-only refinement. The FSC of full reconstructions comparing the two methods shows their agreement far exceeds the recoverable signal (grey). (B) Partial snapshots of the final reconstruction following post-processing, superimposed on PDB ID 3J79 (Wong et al., 2014)*

#### Disk & memory considerations

While GPUs offer high capacity and throughput for data processing, the available memory on the device is limited, which leads to some challenges for its use and management. RELION typically requires large amounts of memory. Fortunately, its peak use is *not* during the accelerated, computationally most intensive parts of the algorithm. Rather, memory usage peaks during the reconstruction step, which is executed on the CPU as described above. The available on-card GPU memory however remains a limitation, as it determines the capacity for storage of the oversampled Fourier transforms of one or more references. This is of particular concern for larger and higher resolution structures, which require more memory to be faithfully represented. When resolving detail at the Nyquist frequency, due to twofold oversampling, we need memory corresponding to twice the image dimension cubed. For example, when using 400^2^-pixel particle images, the required grid size is 800^3^, which becomes ~2GB per class, since each value requires 4 bytes in single precision. Moreover, as the reconstructed object also needs to be accommodated, this number is effectively multiplied by 2.5.

Peak memory usage for particle image sizes up to 400^2^ indicate that at most 6 GB of on-card GPU memory is needed to perform refinement to Nyquist (Fig. 5). The 3D classification is usually performed at much lower resolution, and for this reason the memory requirements are lower.

To enable efficient evaluation and good scaling on GPUs, several new methods to manually manage data at low cost have been implemented. Lower levels of parallelism are coalesced into larger objects using customised tools, which results in more efficient use of memory. In addition, because of the much improved performance, multiple tasks have become limited by how fast input data can be read from disk. Therefore, we now find it highly beneficial to cache data on local solid state devices (SSDs), as has also been observed for GPU-accelerated CTF estimation (Zhang, 2016).

**Figure 5:**
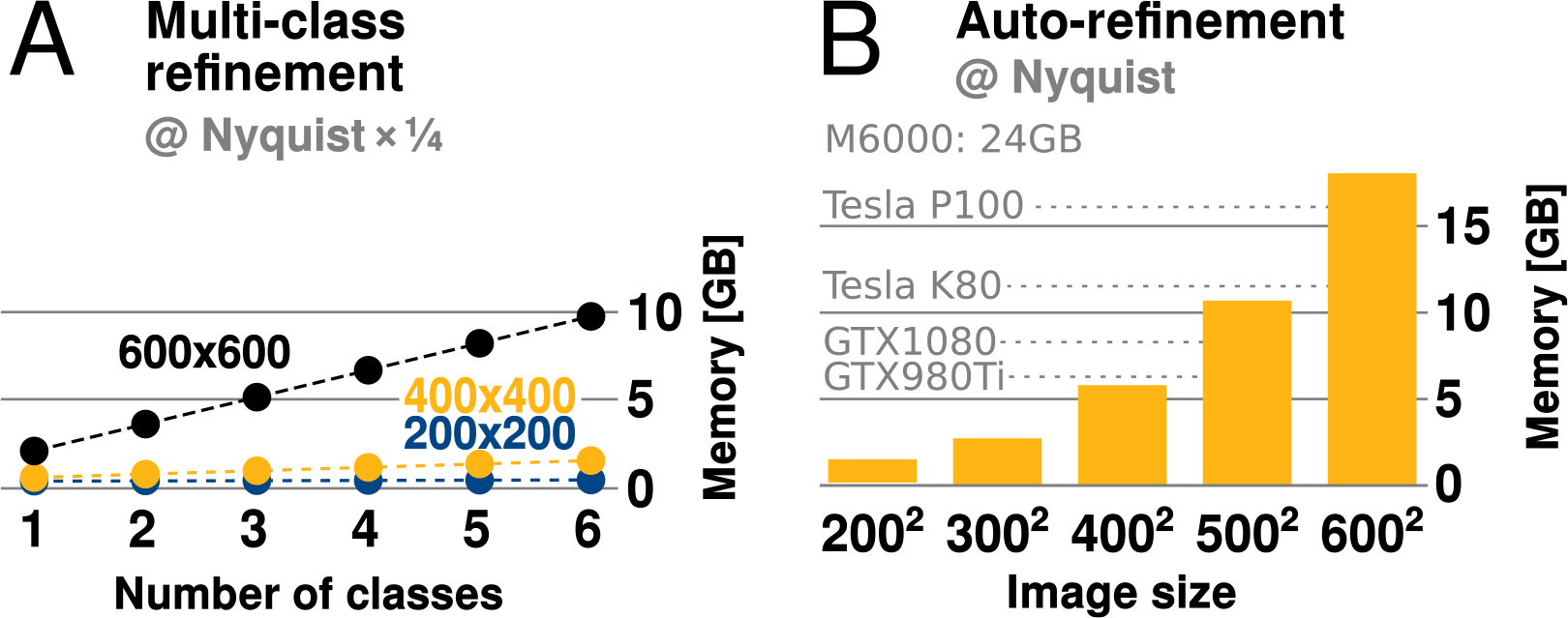
*GPU memory requirements. (A) The required GPU memory scales linearly with the number of classes. (B) The maximum required GPU memory occurs for single-class refinement to the Nyquist frequency, which increases rapidly with the image size. Horizontal grey lines indicate avaliable GPU memory on different cards.*

To allow this in a straightforward way, RELION-2 features the ability to automatically copy data sets to fast local disks prior to refinement. Depending on the storage system used, this can further increase performance during less computationally intensive refinements, such as 2D classification.

### 2.2 Acceleration of automatic particle picking

#### Parallelism in autopicking

RELION implements a template-based particle selection procedure, which is implemented by calculating a probability measure (the R-value) for each pixel in the micrograph to signify the likelihood that it is the location of any of the provided templates (Scheres, 2015). The R-value map of a micrograph considers all possible rotations of each template, and is subsequently used in a peak-search algorithm that locates particles within the original micrograph. These calculations are performed in the Fourier space, where they are extremely efficient (Roseman, 2003). In fact, they are so fast that their execution time becomes negligible compared to the time spent transforming image objects from real space and back using FFTs. Consequently, even though reference templates are also treated as independent tasks to increase parallelism in the GPU version, a much larger gain is found at the level of template rotations. By parallelising this step we have been able to improve the efficiency considerably through parallel execution of FFTs. For example, when using 5-degree incremental template rotations, 72 such inverse FFTs are now performed concurrently on the GPU, through the CUFFT CUDA library. The size of these FFTs is now also padded automatically, since severe performance penalties can occur if the transform size includes any large prime factors.

**Figure 6:**
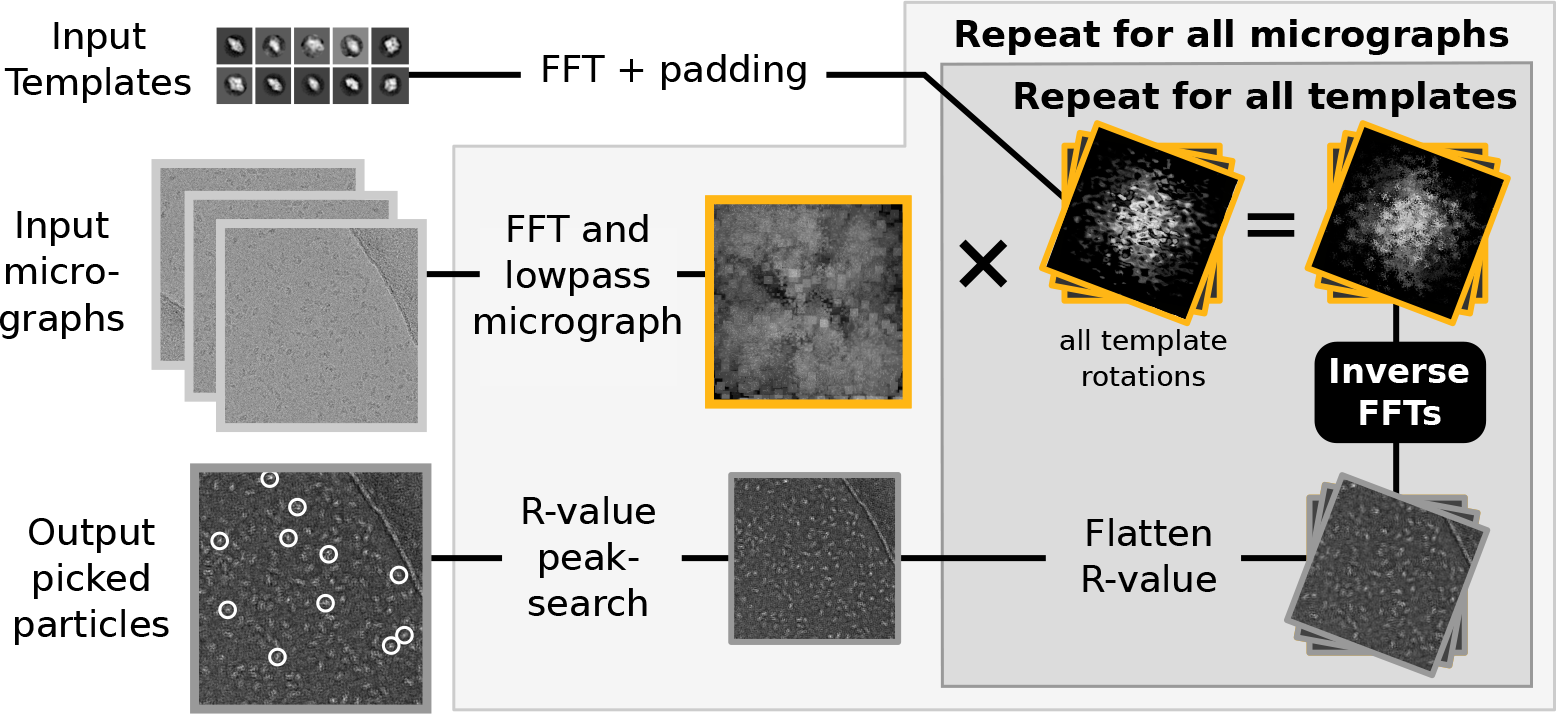
*Semi-automated particle picking in* RELION-*2. The low-pass filter applied to micrographs is a novel feature in* RELION, *and drastically reduces the size and execution time of the highlighted inverse FFTs, which accounts for most of the computational work. In addition to the inverse FFTs, all template-and rotation-dependent parallel steps have also been accelerated on GPUs.*

#### Low-pass filtering of micrographs

Even after parallelisation and acceleration, cross-correlation-based particle selection is still dominated by computing many large inverse FFTs (Fig. 6), as has been observed previously (Castaño-Díez et al., 2008). Reducing their size is thus the most straightforward way to further reduce execution time. Reference templates are typically subject to low-pass filtering, and for this reason we investigated the possibility to apply a similar filtering to all micrographs, which reduces high-frequency information.

We found little difference in the particles selected when discarding resolution information in micrographs beyond that of search templates. While intuitively straightforward, this conclusion drastically reduces the size of FFT grids and subsequent computations, which provides large acceleration at virtually no quality loss. The low-pass filtering also significantly reduces the amount of memory required for particle selection, which permits parallelism to target hardware like desktop workstations much more efficiently.

**Figure 7:**
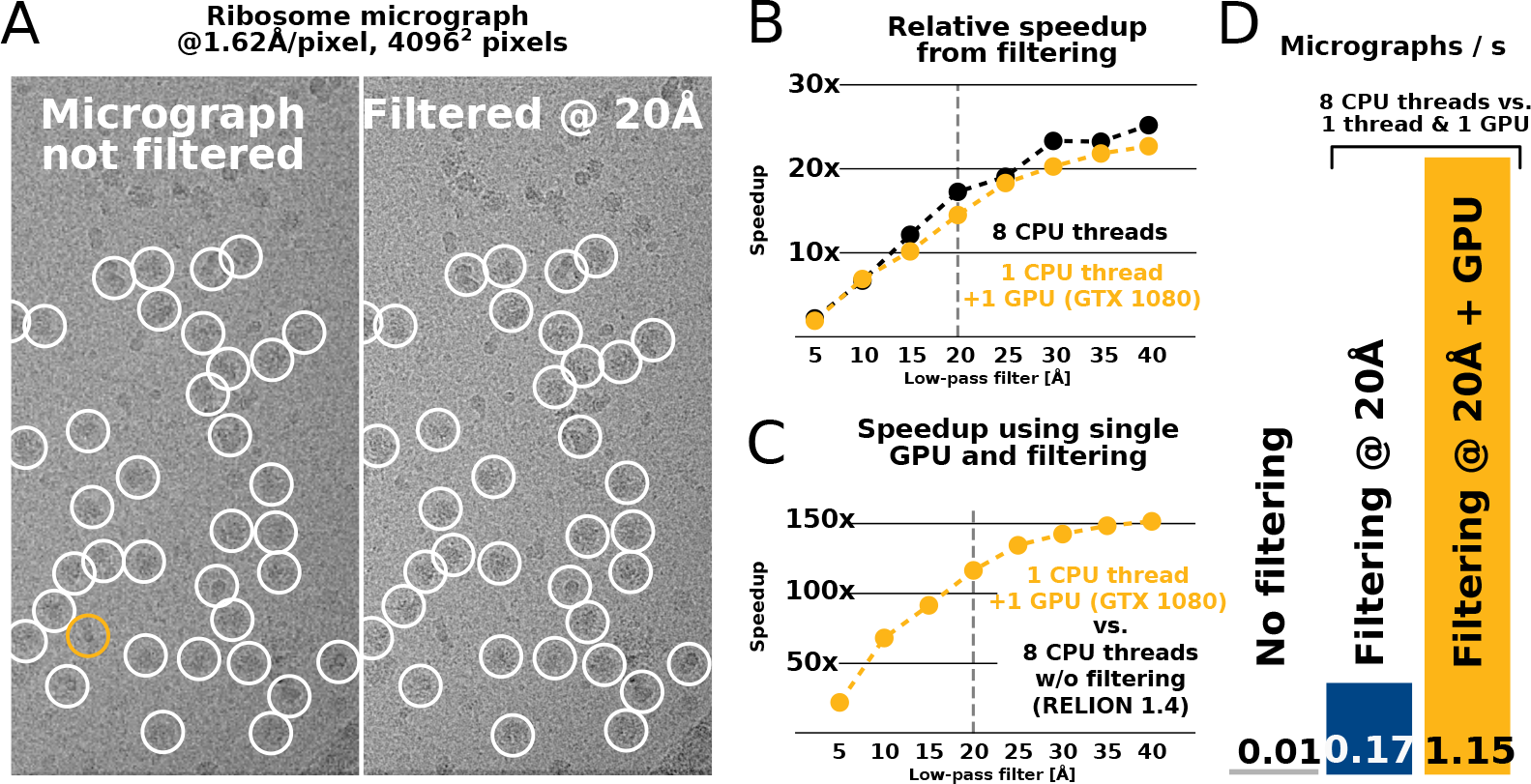
*Low-pass filtering and acceleration of particle picking. (A) Ribosomal particles were auto-picked from representative* 4096^2^ *-pixel micrographs collected at 1.62A/pixel using four template classes, showing near-identical picking with and without low-pass filtering to 20^Å^. The only differing particle is shown in orange, and it likely does not depict a ribosomal particle. (B-C) Despite near-identical particle selection, performance is dramatically improved. (D) Filtering alone provides almost 20-fold performance improvement on any hardware compared to* RELION-*1.4, and when combined with GPU-accelerated particle picking the resulting performance gain is more than two orders of magnitude using only a single GPU.*

#### Autopicking performance

We tested both the speed and the quality of picked particles of our new implementation. In an initial test, a single 4096^2^-pixel micrograph containing ribosomes at 1.62 Å/pixel was processed against 8 templates with 5 degree angular sampling and no low-pass filtering. This took 675s to evaluate on a CPU-only workstation (i7-5960X, using 1 thread merely for reference). When applying low-pass filtering to 20Å, this time is reduced to 39s, i.e. by a factor ~17. When using a single consumer-level GPU (GTX1080) in a single CPU thread, execution is further reduced to just 0.73s, i.e. an additional factor ~54. A workstation with 8 CPU threads and a single GPU can therefore now process ~925 micrographs in the same time previously required to process just 8 micrographs (1 per available core) as shown in Fig. 7.

We further evaluated the quality of filtered selection according to the *β*-galactosidase benchmark (EMPIAR entry 10017) used in the original implementation in RELION-1.3 (Scheres, 2015). This data set consists of 84 micrographs of 4096^2^ pixels (1.77 Å/pixel), and comes with coordinates for 40,863 particles that were manually selected by Richard Henderson. The latter were used for comparison with our autopicking results, with a center cutoff distance of 35 pixels for particles to be considered identical (Tab. 1). Filtered selection did not decrease the quality of the results, but rather provided an increased recall without increasing the false discovery rate (FDR, see e.g. Langlois and Frank (2011) for definitions of recall and FDR). When filtering and GPU-acceleration are combined, a single GPU provides almost three orders of magnitude performance improvement over a single CPU core.

**Table 1:**
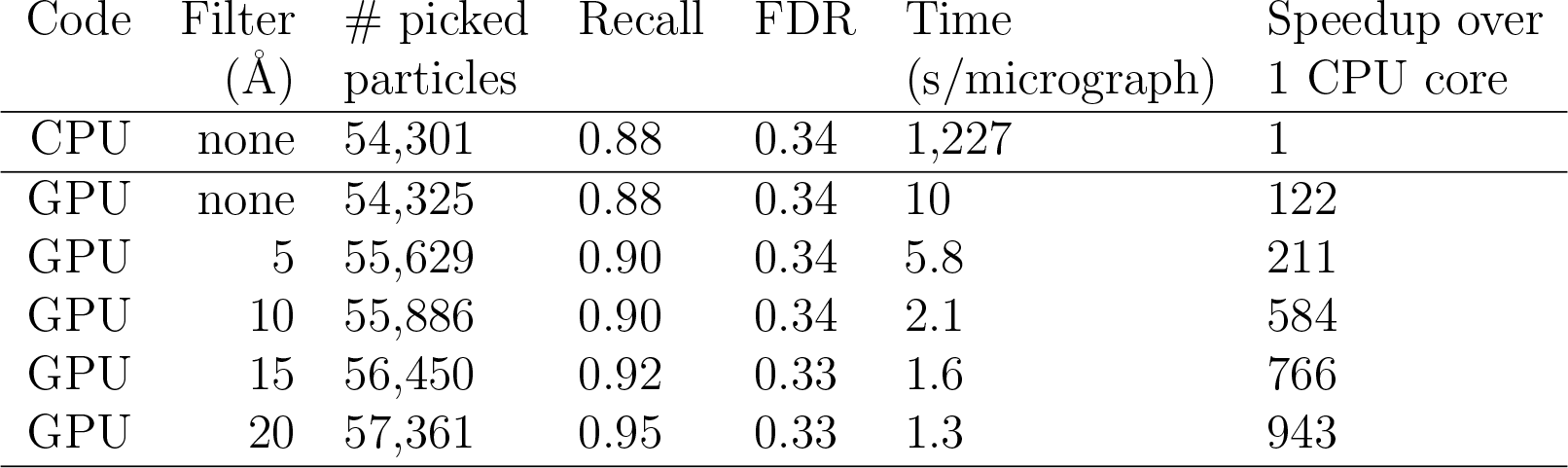
Quality and speed of autopicking for the *β*-galactosidase benchmark. Comparing the CPU version with the GPU version using increasing levels of low-pass filtering yields progressively higher recalls at similar FDRs. The GPU version yields identical results to that of the CPU version, but at a much reduced computational costs. Filtering does not depend on GPU-acceleration, and will perform similarly using only CPUs.

| Code | Filter<br>(Å) | # picked<br>particles | Recall | FDR | Time<br>(s/micrograph) | Speedup over<br>1 CPU core |
| --- | --- | --- | --- | --- | --- | --- |
| CPU | none | 54,301 | 0.88 | 0.34 | 1,227 | 1 |
| GPU | none | 54,325 | 0.88 | 0.34 | 10 | 122 |
| GPU | 5 | 55,629 | 0.90 | 0.34 | 5.8 | 211 |
| GPU | 10 | 55,886 | 0.90 | 0.34 | 2.1 | 584 |
| GPU | 15 | 56,450 | 0.92 | 0.33 | 1.6 | 766 |
| GPU | 20 | 57,361 | 0.95 | 0.33 | 1.3 | 943 |

### 2.3 A complete workflow for *β*-galactosidase

To illustrate the impact of our GPU implementation and show how it can alter practical work, we chose to re-analyse the EMPIAR-10061 dataset of *β*-galactosidase (Bartesaghi et al., 2015) using relion-2. Downloading this 12.4 TB set took a total of 151h (6.3 days) using the ascp client for fast file transfer. We performed the entire processing workflow, including initial beam-induced motion correction in UNBLUR (Grant and Grigorieff, 2015), CTF estimation in Gctf (Zhang, 2016), automated particle picking, 2D and 3D classification, movie-refinement, particle polishing (Scheres, 2014) and high-resolution auto-refinement, on a single workstation with four GTX 1080 cards. Calculating a map to 2.2 Å resolution (Fig. 8A) took under five days - less time than downloading the data. Figure 8B shows an overview of the most computationally demanding steps. The parts of the workflow that have been GPU-accelerated no longer dominate execution, but this exposes other new bottlenecks. In particular, steps that involve reading large movie files from disk become a problem. We also note that due to the extremely rigid nature of the *β*-galactosidase complex, only a single 3D classification step was performed. This is unrealistic for many other structures: typically 3D classification is repeated multiple times. In such a scenario, the impact of the GPU acceleration is even larger. Even when repeating 3D classification five times using a workstation for less than $10,000, processing would still be faster than downloading the data.

**Figure 8:**
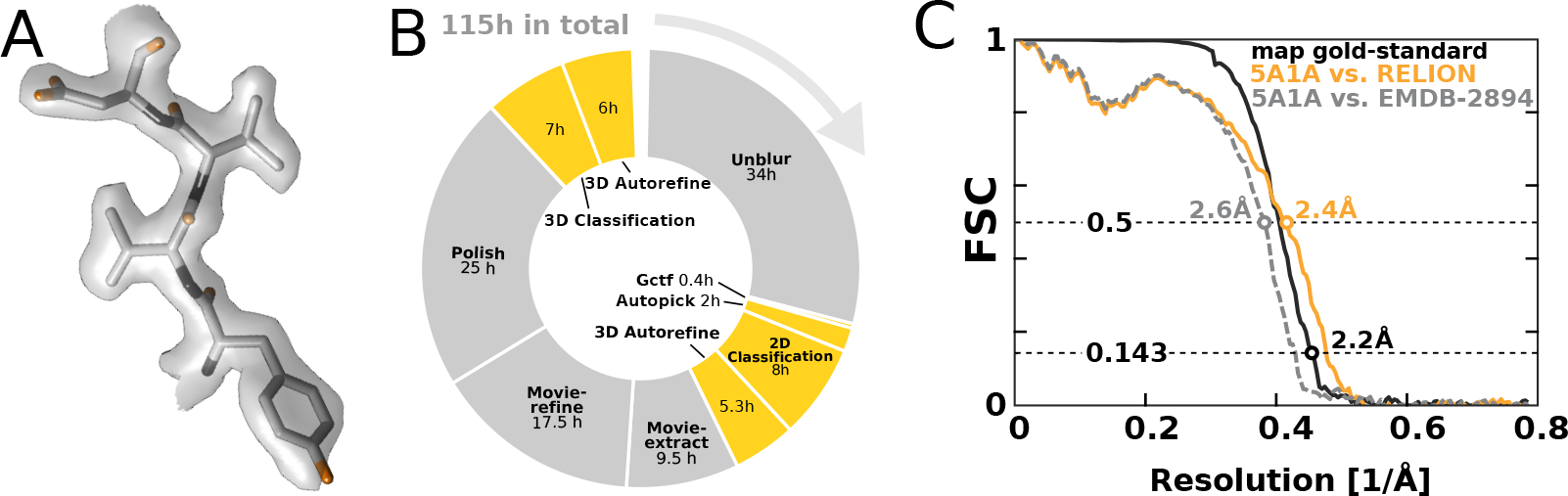
*High-resolution structure determination on a single desktop machine. (A) The resulting 2.2 Å map shows excellent high-resolution density throughout the complex. (B) The most time-consuming steps in the image processing workflow. GPU-accelerated steps are indicated in orange. The total time of image processing was less than that of downloading the data. (C) The resolution estimate is based on the gold-standard FSC after correcting for the convolution effects of a soft solvent mask (black). The FSC between the* RELION *map and the atomic model in PDB ID 5A1A is shown in orange. The FSC between EMDB-2894 and the same atomic model is shown for comparison (dashed gray).*

## 3 Discussion

We present a GPU-enabled implementation of RElion-2, as a first step to address current and future needs for large and expedient computations in the field of cryo-EM structure determination. The principal benefits drawn from the presented work are twofold. First, the nature of progress in scientific applications is to continually re-evaluate and examine data in many different ways. With ease of re-processing data, the threshold for trial, error and successive improvement of existing methods is now markedly lowered. Second, the order-of-magnitude speedups make it possible to get by with much less hardware for cryo-EM processing, in most cases even desktops. This removes a computational bottleneck for large labs, and enables any group to perform their own reconstruction without access to supercomputers.

In the next few years, larger data sets and image sizes are expected, as well as new methods that require expedient processing of large data sets. The large reduction in computational costs opens up the possibility to perform more ambitious computational analyses without increasing the investments. For example, the favourable scaling of performance we observed for multiclass refinements will make it feasible to use many more classes than was practical before, which will lead to better descriptions of conformational diversity in flexible molecules. Additionally, with even faster algorithms and hardware it might soon be possible to perform highly automated, on-the-fly, structure determination while data acquisition is ongoing. In anticipation of these developments, relion-2 already implements a pipelined approach for automated execution of pre-determined image processing workflows (details to be published elsewhere).

While the new GPU implementation has removed many of the previous computational bottlenecks in RELION, the large speedup has exposed several new areas of the code that can now dominate execution time, such as data input/output and the reconstruction step during iterative image refinement. Although these parts of the algorithm were previously insignificant, in some cases they now collectively account for roughly 50% of total execution time. These parts of the code will see benefit from further modifications. Future work will e.g. strive to further generalise parallelism such that performance is less dependent on the type of refinement performed, as sufficient parallelism is always available within the RELION core algorithm. Memory requirements on the GPU are also expected to be reduced further, so that larger image sizes and more classes can be handled to higher resolution.

With the current implementation, cryo-EM structures to near-atomic resolution can be calculated in a matter of days on a single workstation, or hours on a GPU-cluster. Nevertheless, the aim of the current adaptations is not to present a final solution to computational needs in relion; while the present version achieves excellent speedup on a wide range of low-cost systems, we expect the acceleration to improve both in performance and coverage. Generalising the low-level parallelism described here to vectorised CPU calculations, and possibly an open GPU language like OpenCL, will constitute little more than translating this parallelism to new instructions. This is something we intend to pursue in the future. As such, relion-2 represents a new incarnation of an existing algorithm, which is intended to be developed far further in the following years. Meanwhile, we hope that the current implementation will have as much impact in the broader community as it is already having in our labs.

## 4 Materials and Methods

### Data sets & hardware specifications

The ribosome data used for the 3D classification and refinement in Figs. 3 and 4 correspond to EMPIAR entry 10028 (Wong et al., 2014). For autopicking, EMPIAR entry 10017 was used (Scheres, 2015). The complete workflow for *β*-galactosidase used EMPIAR entry 10061 (Bartesaghi et al., 2015). The workstation used for ribosome refinement and autopicking was equipped with a single Intel Core i7-5960X CPU with 8 cores running at 3GHz and 64GB memory. The corresponding CPU cluster calculations used 10 compute nodes equipped with dual Xeon E5-2667 or E5-2643 CPUs (12 physical cores, for a total of 120 cores) running at 2.9-3.4GHz with 64GB memory. Refinement tests were run with four consumer GPUs (testing both NVIDIA GTX 980Ti and GTX 1080 cards), and two CPU threads per GPU to improve utilisation. The classification speedup (Fig. 3C) compares the GPU workstation to a single cluster node, which is hardware of roughly comparable price (a modern E5-2643v4 node might be 10% cheaper). The autopicking benchmark only used a single GPU to better reflect a simple desktop setup. Admittedly, comparing performance to a single core is not entirely fair when a typical CPU has multiple cores, but in practice a workstation can easily be equipped with multiple GPUs too. The results for the *β*-galactosidase test case were obtained on a single desktop machine with four GTX 1080 cards, a 400GB local SSD scratch disk, dual Xeon E5-2620 CPUs (12 cores in total) running at 2.4GHz and with 64GB memory.

### *β*-galactosidase image processing

Super-resolution 8*k* × 8*k* micrograph movies with 38 frames were submitted to initial beam-induced motion correction using UNBLUR (Grant and Grigorieff, 2015). The resulting average micrographs were used for CTF estimation in Gctf (Zhang, 2016). Autopicking with six templates yielded an initial data set of 130,375 particles, which were subjected to reference-free 2D classification using 200 classes. This initial classification was done using 4× downscaled particles (with a pixel size of 1.274 Å and a box size of 192 pixels). Selection of the 75 best classes resulted in 120,514 particles. All subsequent calculations were performed using 2× downscaling (resulting in a pixel size of 0.637 Å and a box size of 384 pixels). The selected particles were subjected to an initial 3D auto-refinement that used PDB ID 3I3E (Dugdale et al., 2010) as an initial model. Subsequent movie-refinement (with a running average of 7 movie frames and a standard deviation of 2 pixels on the translations) was followed by particle polishing (using a standard deviation of 1000 pixels on the inter-particle distance). The resulting shiny particles were submitted to a single round of 3D classification with exhaustive 7.5- degree angular searches and eight classes. Selection of the seven best classes yielded a final data set of 109,963 particles, which were submitted to 3D autorefinement. The final resolution was estimated using phase-randomisation to account for the convolution effects of a solvent mask on the FSC between the two independently refined half-maps (Chen et al., 2013). This mask was generated by binarisation of a 15 Å low-pass filtered version of the reconstructed map, with addition of a five-pixel wide cosine-shaped soft edge. FSC curves between the model and the solvent-masked map were calculated with `relion_image_handler`. The same soft solvent mask was also used for the calculation between EMDB-2984 and the atomic model.

## 5 Acknowledgements

The authors would like to thank Szilárd Páll, Nikolay Markovskiy, and Mark Berger for fruitful discussions and CUDA suggestions; Shintaro Aibara and Marta Carroni for providing data and early quality testing; and Toby Darling, Jake Grimmett, and Stefan Fleichmann for technical support.

## 6 Competing interests

SHWS: Reviewing editor, eLife.

The other authors declare that no competing interests exist.

## 7 Funding information

Medical Research Council (MC_UP_A025_1013): Sjors HW Scheres

Swedish Research Council (2013-5901): Erik Lindahl

Swedish e-Science Research Centre: Erik Lindahl

BioExcel CoE (EINFRA-2015-1-675728): Erik Lindahl

The funders had no role in study design, data collection and interpretation, or the decision to submit the work for publication.

